# Estimating changes in RNA Polymerase II elongation and intron splicing speeds from total RNA-seq data

**DOI:** 10.1101/2025.08.24.672013

**Authors:** Jakub Koubele, Andreas Beyer

## Abstract

We present a computational method to jointly estimate changes in the elongation speed of RNA Polymerase II, and the speed of intron splicing, using total RNA-seq. An implementation of our method is available at https://github.com/jkoubele/pol-ii-speed.

## 1 Introduction

The speed of RNA polymerase II (Pol II), measured in the number of nucleotides synthesized per unit of time, plays an important role in RNA metabolism. Changes in Pol II speed can affect several co-transcriptional processes, such as splicing, m^6^A RNA methylation, and polyadenylation (Muniz et al., 2021). Recently, a study of Debès et al. (2023) showed that organism aging is associated with an increase in Pol II speed, whereas lifespan-prolonging interventions tend to decrease it.

The speed of intron splicing (measured in time needed to splice the intron out of the transcript) is another important aspect of RNA metabolism. Splicing time can vary greatly, ranging from several seconds up to about 15 minutes (Shenasa and Bentley, 2023).

Measuring the Pol II elongation speed or the speed of splicing is a challenging task, and a variety of biochemical methods were used for that purpose.

To overcome the necessity of specialized biochemical experiments, it was proposed that the changes in Pol II speed can also be estimated from total RNA-seq (Ameur et al., 2011; Jonkers et al., 2014; Kawamura et al., 2019; Debès et al., 2023). Total RNA-seq (specifically Ribo-minus RNA-seq) is a widely used protocol, and many such datasets are publicly available.

Total RNA-seq consists of both the nascent and mature RNA. Changes in gene expression affect both the nascent and mature components, while changes in Pol II speed and splicing time affect the nascent RNA only. As introns are predominantly present in the nascent RNA (with the exception of intron retention events), changes in intron coverage can be linked to changes in Pol II speed and splicing time.

The previous methods mentioned above used the gradient of intron coverage for the estimation of Pol II speed. By contrast, exonic coverage is usually dominated by mature RNA, with only a small fraction of reads originating from nascent RNA, making it less suitable for estimating changes in elongation or splicing speeds.

Here, we present a novel computational method that jointly estimates changes both in Pol II speed and splicing speed from total RNA-seq.

## 2 Model derivation

We will now derive our model, which works with samples of total RNA-seq. For notation, we will be using **bold** symbols to denote vectors, and normal font to denote scalars.

### 2.1 Sample annotation

The data consists of total RNA-seq samples. Each sample is annotated by a vector of explanatory variables **x**. The explanatory variables may consist of categorical factors, such as genotype or intervention, encoded by one-hot encoding. The continuous variables, e.g., age measured in weeks, may also be present in the **x**. The intercept element (constant 1) is not supposed to be included in the **x** (we will introduce intercept terms as separate parameters later on).

Each sample also has some library size factor *s*, representing the sequencing depth. It may be given simply as the number of reads; in our implementation, we decided to estimate *s* by the median-of-ratios method described in (Anders and Huber, 2010). We will also work with the logarithm of the library size, *𝓁* = log(*s*).

### 2.2 Modeled gene and intron properties

In each gene, we will model three properties of interest: the gene expression rate, the speed of Pol II elongation, and the time needed for intron splicing.

The gene expression rate *η* is modeled as

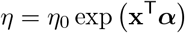

The unit of *η* is the amount of polymerases starting transcription per second, [*η*] = polymerases · s^−1^. The vector ***α*** is a model parameter, containing the log-fold change (LFC) effect of the explanatory variables. The term *η*_0_ is an intercept term of the gene expression, with the same unit as *η*.

The remaining two properties, speed of Pol II elongation and time needed for intron splicing, are defined for each intron in the gene separately.

The Pol II speed *v* is modeled as

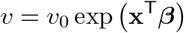

The unit of *v* is the amount of base-pairs transcribed per second, [*v*] = bp · s^−1^. Similarly as above, ***β*** is a vector of LFC effect of explanatory variables on Pol II speed, and *v*_0_ is an intercept term with the same unit as *v*.

For each intron, we also model the time needed to splice it out (and degrade it) as

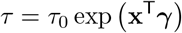

This splicing time is measured in seconds, [*τ*] = s. As above, ***γ*** is a vector of LFC effects of explanatory variables, and *τ*_0_ is an intercept, also measured in seconds.

Note that *τ* ^−1^ is the speed of splicing, and −***γ*** is the LFC effect of explanatory variables on the splicing speed; we simply decided to work with the splicing time rather than splicing speed in our notation.

### 2.3 Intron coverage distribution

The central part of our model is built on the fact that all gene properties introduced above - expression rate *η*, Pol II speed *v*, and splicing time *τ* - affect the read coverage of introns. We will therefore use the intron coverage (together with the count of exonic reads) to estimate the LFC vectors ***α, β***, and ***γ***.

The relation between the Pol II speed and intron coverage was first described by (Ameur et al., 2011), and used by subsequent work of others.

The read coverage of any genomic region (intron in our case) can be decomposed into two parts: the location of the reads, and the number of reads observed. The location component can be described by a probability distribution over the intron interval, so the locations of individual reads can be understood as samples from such a distribution. The number of reads observed is also a random variable, specified by some count distribution.

We will show that the location of intronic reads depends on the Pol II speed *v* and the splicing time *τ*, but not on the expression rate *η*. However, the amount of reads observed in the intron depends on all modeled variables: *η, v*, and *τ*.

#### 2.3.1 Transcribing and unspliced introns

The intronic reads we observe may come from two distinct sources: either from the introns that are currently being transcribed by Pol II, or from the introns that were already fully transcribed, but are not spliced and degraded yet (see Figure 1).

**Figure 1:**
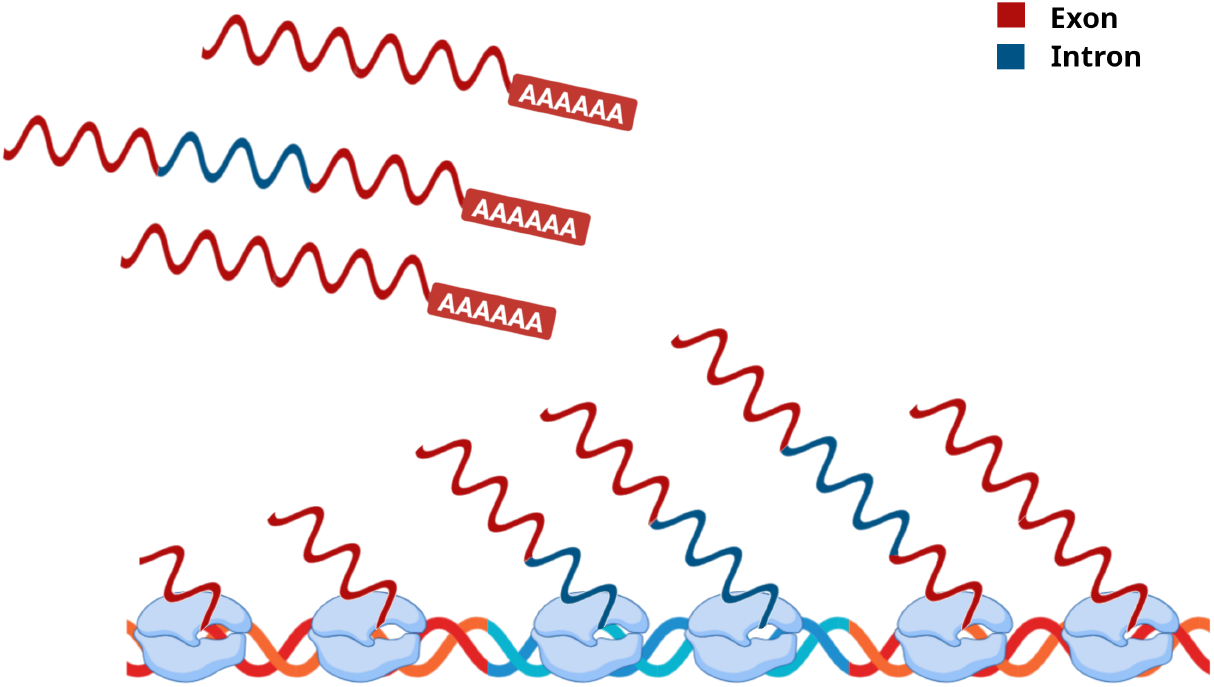
Intronic RNA (in blue) in total RNA-seq may come from two distinct sources: either the intron is currently being transcribed by RNA polymerase, or it was already finished, but not spliced and degraded yet. The finished introns may still be part of the nascent RNA or may come from retained introns in mature RNA.

While we are not able to directly distinguish the source of a given sequencing read, we still can estimate the proportions of reads from these two sources in our data, using the observed intron coverage, as described later on.

Consider an intron of a length *L*; the unit of length is base-pairs, [*L*] = bp. The number of observed reads coming from currently transcribed introns is proportional to the amount of polymerases transcribing the intron:

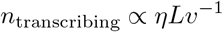

Note that *Lv*^−1^ is the time needed to transcribe the intron. By plugging in the formulas for *η* and *v* from their definitions, we get

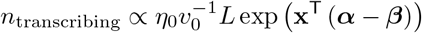

The amount of transcribing introns is therefore inversely proportional to the Pol II speed. With faster Pol II, there will be less Pol II transcribing the intron, resulting in a lower number of such intronic reads; see the illustration in Figure 2.

**Figure 2:**
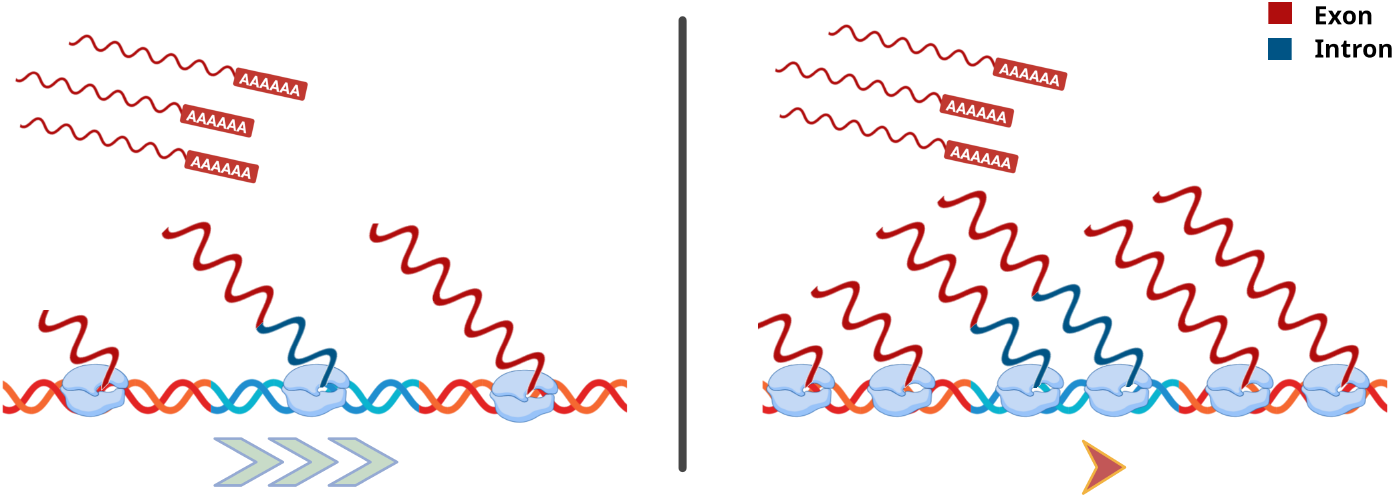
Effect of the change in Pol II speed. In the condition with faster polymerase (left), there is less polymerases currently transcribing a gene, compared to the condition with slower polymerase (right). Note that the amount of mature RNA is not affected by the change in elongation speed.

The number of reads from introns that were fully transcribed, but not spliced out and degraded yet, is proportional to the expression rate and the splicing time:

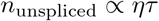

By plugging in the formulas for *η* and *τ*, we get

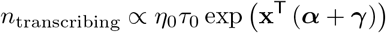

#### 2.3.2 Intron coverage as a mixture distribution

The two kinds of intronic reads (either currently transcribing or finished ones) described above have a different distribution of their location in the intron.

For notational simplicity, we will assume the intron to be a continuous interval [0, 1], oriented in the 5’ to 3’ direction. The location of any read is then a real number on this interval.

The fully finished introns can produce reads anywhere in the intron. Therefore, the location of these reads follows a uniform distribution, with a density

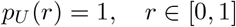

On the other hand, the transcribing introns can produce reads only in the part of the intron that was already transcribed. Due to this, the location of these reads follows a distribution with decreasing density. In our implementation, we will work with a simple triangular distribution, with the density function

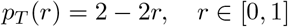

The overall distribution of intronic reads is then a mixture of the uniform and triangular distributions, with a density

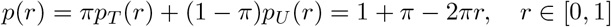

The variable *π* ∈ [0, 1] is the mixture weight, giving the proportion of the triangular density; the term (1 − *π*) is then the proportion of the uniform distribution. The density is visualized in Figure 3.

**Figure 3:**
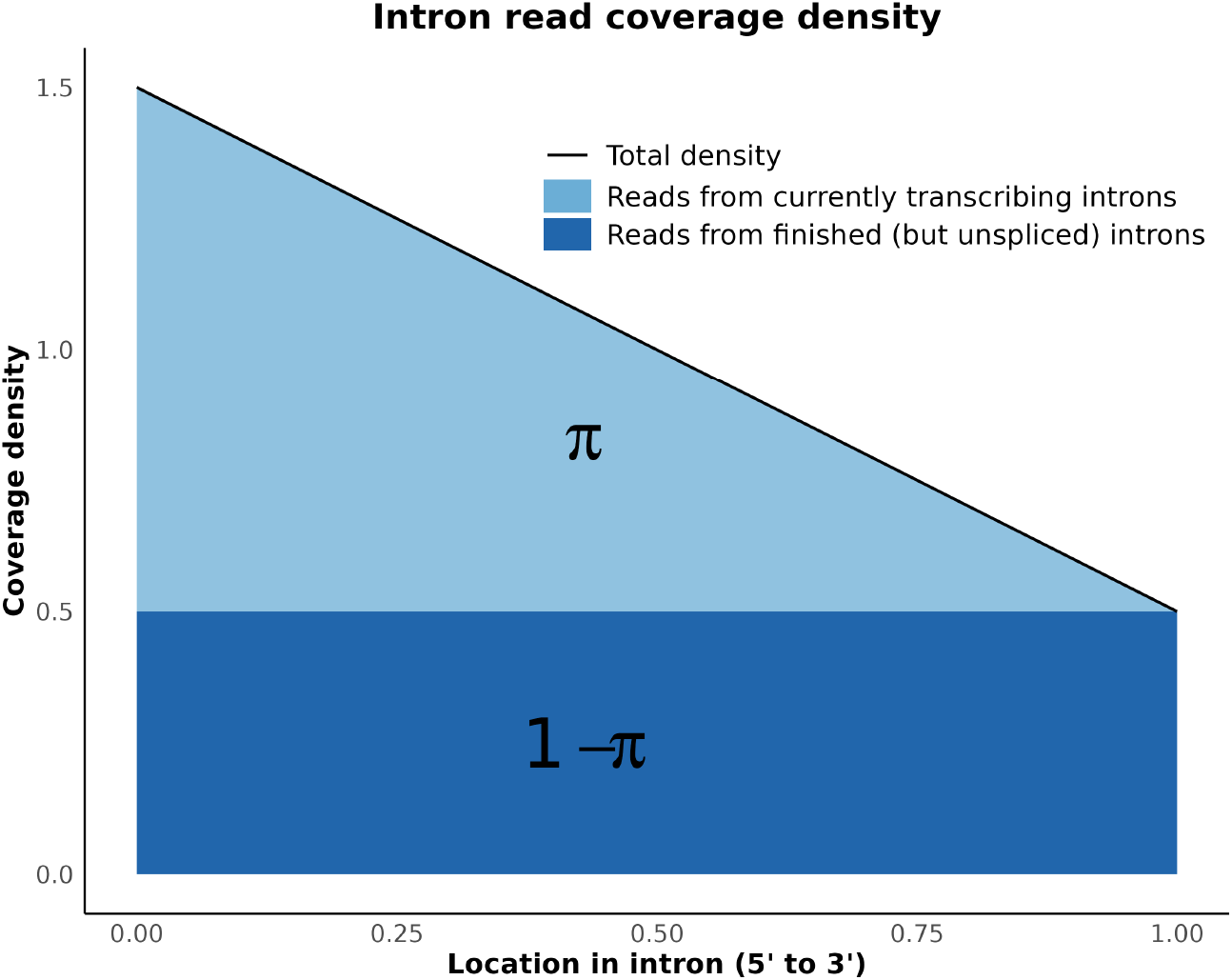
Density function of the intronic read location. The resulting distribution is a mixture of a decreasing triangular distribution, with a weight *π*, and a uniform distribution, with a weight (1 − *π*).

The variable *π* is not a free parameter in our model. Instead, it depends on the number of reads from transcribing and unspliced introns, as

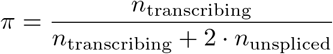

The factor 2 in front of *n*_unspliced_ accounts for the fact that fully transcribed introns are on average 2× longer than currently transcribing introns, and therefore produce 2× more reads.

We will now plug in the formulas for *n*_transcribing_ and *n*_unspliced_, obtaining:

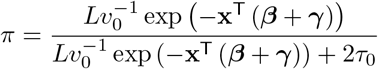

We will now denote

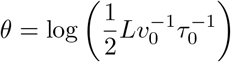

We can then simplify the formula for *π* as

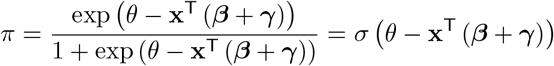

where *σ*(.) denotes the logistic (sigmoid) function,

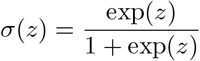

The variable *θ* will be a parameter in our model. Note that the term 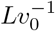 is an intercept time needed to transcribe the intron, and *τ*_0_ is an intercept time needed to splice the intron out. The term 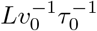, which appears in the definition of *θ*, is therefore an intercept fraction of transcription and splicing times.

### 2.4 Exon and intron read counts

Besides the location of intronic reads, we will also be using the count of exonic and intronic reads to fit our model.

In our implementation, we align all reads to the genome at first. Then, we evaluate which reads have sufficient overlap with some intron, and we construct an intron count matrix from these reads. After that, we take all reads that didn’t overlap with any intron, and we align them to the transcriptome. This way, we can account for different transcript lengths stemming from an alternative splicing of exons.

#### 2.4.1 Exon read counts

In our implementation, we are aligning exonic reads (i.e., those without sufficient overlap with any intron) to the transcriptome, obtaining a count of exonic reads for every gene and sample. We are modeling this observed count as a sample from some count distribution; we used the Poisson distribution in our implementation for simplicity, although other distributions can also be used (e.g., the negative binomial distribution).

We model the mean of the Poisson distribution as

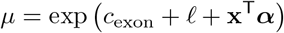

The term *c*_exon_ is an intercept term for the given gene (and is therefore the same for all samples). The term *𝓁* is the (sample-specific) logarithm of the library size, introduced above.

We will now discuss two caveats of our model. The first caveat is that we model the *µ* to depend on the LFC of gene expression ***α***, but not on the LFC of Pol II speed ***β***. However, a small fraction of exonic reads comes from the nascent RNA (see Figure 1), the abundance of which depends on the Pol II speed. Here, we assume that this fraction is reasonably small, so the majority of exonic reads come from the mature RNA, and we are omitting the link between *µ* and ***β***.

The second caveat comes from the fact that the abundance of mature RNA may also be regulated by its degradation rate, not only by its expression. While the transcription initiation is usually under much stricter regulation than the RNA degradation, there are known examples of genes that are strongly regulated by control of their degradation rate, e.g., many mRNA important in the immune system (Schott and Stoecklin, 2010). In such cases, our model will incorrectly attribute the changes in exon read counts to the changes in the gene expression rate *η*, which will consequently compromise the estimated values of all model parameters.

#### 2.4.2 Intron read counts

The observed intron read counts are also samples from some count distribution, which we model by a Poisson distribution in our implementation. Contrary to exonic reads, we are modeling the read count of every intron in the gene separately.

We will now derive the formula for the mean of the Poisson distribution for intronic reads, denoted as *ν*. In the given sample, the value of *ν* depends on the number of transcribing and unspliced introns as

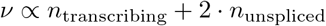

As above, the factor of 2 in front of *n*_unspliced_ accounts for the different average length of transcribing and unspliced introns.

We will now plug-in the formulas for *n*_transcribing_ and *n*_unspliced_ into the formula for *ν*. We obtain:

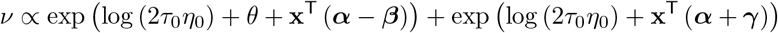

We will now introduce the intercept term *c*_intron_ and account for the library size of the sample. This will get us:

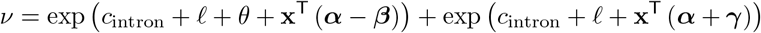

The term log (2*τ*_0_*η*_0_) is an unknown constant and was absorbed into the intercept *c*_intron_.

### 2.5 Model summary

We will now summarize the model derived above.

#### Sample annotation

Each sample is associated with:

- Vector of explanatory variables **x**.
- Logarithm of library size *𝓁*.

#### Model parameters

For each gene, the model has these parameters:

- Vector ***α***, denoting the LFC effects (of explanatory variables) on gene expression.
- Intercept term *c*_exon_.

The following parameters are specified for each intron:

- Vector ***β***, denoting LFC effects on Pol II speed.
- Vector ***γ***, denoting LFC effects on splicing time.
- Intercept term *c*_intron_.
- Parameter *θ*, related to the intercept ratio of transcription and splicing times.

#### Model output

Given the sample annotation and model parameters, the model predicts following values for each sample.

- Value *µ*, a mean of Poisson distribution for exon read count. It is specified by the formula

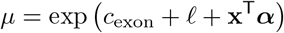

The following values are predicted for each intron separately:

- Value *ν*, a mean of Poisson distribution for intron read count, given as:

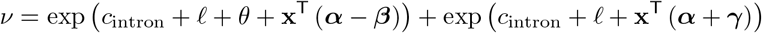
- Value *π*, a mixture weight of a triangular distribution in the intron coverage density, computed as:

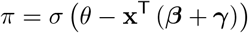

Given the value of *π*, the density of intron coverage is then:

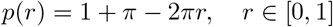

where *r* is the relative position in the intron, oriented in the 5’ to 3’ direction and normalized into a [0, 1] interval.

## 3 Model fitting and inference

The model is trained by maximum likelihood estimation (MLE). The loss function (negative log-likelihood) is a sum of 3 components: the loss for exon read counts prediction (one value per gene), the loss for intron read counts prediction (one value for every intron in a gene), and the loss for location of intronic reads (intron coverage), also calculated for every intron.

The loss function for exon and intron read counts is the negative log-likelihood of the Poisson distribution. We will now elaborate on the details of the loss function of intron coverage.

### 3.1 Loss function of intron coverage

For given intron, the coverage loss is given by the negative log-likelihood of the predicted coverage density, that is:

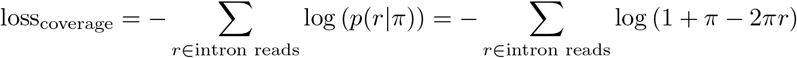

Where *r ∈* [0, 1] is a position of a read within the intron.

In our implementation, we find it useful to discretize the positions of reads into a fixed number of bins, and work with the resulting histogram. This way, we treat all reads in a given bin as having the same location, and we therefore need to loop only over the bins rather than over all reads when evaluating the coverage loss function.

### 3.2 Inference

To assess the significance of the estimated LFC parameters (i.e., the vectors ***α, β***, and ***γ***), we employed a likelihood-ratio test. Specifically, after fitting the full model, we fitted a restricted version in which a given element of one LFC vector was fixed to zero. The difference in the loss functions (negative log-likelihoods) between the restricted and unrestricted models was then used to perform a *χ*^2^ test.

## 4 Implementation

We implemented our model in PyTorch (Ansel et al., 2024), making use of its automatic differentiation system for model fitting. The surrounding workflow, from pre-processing of the input FASTQ data to model fitting, was orchestrated with Nextflow (Di Tommaso et al., 2017).

## 5 Conclusion

We presented a computational framework that jointly estimates changes of RNA Polymerase II elongation speed and intron splicing time from total RNA-seq data. By accounting for both processes in a unified model, our method provides a more comprehensive view of co-transcriptional dynamics and may serve as a tool for studying RNA metabolism in diverse biological contexts.

## Code availability

An implementation of our method is available at https://github.com/jkoubele/pol-ii-speed.

## Acknowledgements

This work was funded by the Deutsche Forschungsgemeinschaft (DFG, German Research Foundation) under Germany’s Excellence Strategy – EXC 2030 – 390661388.

Illustrations were created with BioRender.com.

